# Assessing the *in vitro* resistance development in Enterovirus 71 in the context of combination antiviral treatment

**DOI:** 10.1101/2020.02.10.935536

**Authors:** Kristina Lanko, Chenyan Shi, Shivaprasad Patil, Leen Delang, Jelle Matthijnssens, Carmen Mirabelli, Johan Neyts

**Author notes:** Corresponding author, E-mail address, KU Leuven, Department of Microbiology and Immunology, Rega Institute for Medical Research, Laboratory of Virology and Chemotherapy, Herestraat 49 - box 1043, 3000 Leuven, Belgium.

## Abstract

There are currently no antivirals available to treat infection with enterovirus A71 (EV-A71) or any other enterovirus. The extensively studied capsid binders select rapidly for drug-resistant variants. We here explore whether the combination of two direct-acting enterovirus inhibitors with a different mechanism of action may delay or prevent resistance development to the capsid binders. To that end, the in vitro dynamics of resistance development to the capsid binder pirodavir was studied either alone or in combination with (i) a viral 2C-targeting compound (SMSK_0213), (ii) a viral 3C-protease inhibitor (rupintrivir) or (iii) a viral RNA-dependent RNA polymerase (RdRp) inhibitor [7-deaza-2’*C*-methyladenosine (7DMA)]. We demonstrate that combining pirodavir with either rupintrivir or the nucleoside analogue 7DMA delays the development of resistance to pirodavir and that no resistance to the protease or polymerase inhibitor develops. The combination of pirodavir with the 2C inhibitor results in a double-resistant virus population. The deep sequencing analysis of resistant populations revealed that even though resistant mutations are present in less than 30% of the population, this still provides the resistant phenotype.

## Introduction

Enterovirus 71 (EV-71) belongs to the family of Picornaviridae - small positive stranded RNA viruses, comprising many important human pathogens. EV-71 can cause a wide range of clinical manifestations: from self-resolving hand, foot and mouth disease to life-threatening conditions like meningitis, encephalitis and acute flaccid myelitis ^1^. Together with EV-68, EV-71 is attracting more attention in the last years due to the rising incidence of neurological complications, caused by these infectious agents ^2,3^. EV-71 circulates mostly in the Asia-Pacific region, e.g. China and Thailand ^4,5^. There is no antiviral therapy available. A vaccine for EV-71 based on an inactivated C4 genotype virus has been approved in China and shows good protection; however, it does not protect against infections with all genotypes ^6,7^. Several small molecules proceeded to clinical trials as enterovirus inhibitors. Capsid binders bind to the viral particle in the pocket of structural protein VP1, which is normally occupied by a lipid pocket factor. This interaction stabilizes the capsid and prevents conformational changes necessary for the receptor binding and genome release of the virus ^8^. One of the advantages of capsid binders is their broad-spectrum antiviral activity (e.g. pleconaril inhibits both human rhinovirus and enterovirus species) ^9,10^. However, rapid development of resistance to capsid binders in vitro and in vivo has been reported ^9,11–13^. Natural resistant variants are circulating as well ^12,14^. The capsid binder pocapavir is currently being developed as a tool in the polio endgame ^12^.

The 3C protease inhibitor rupintrivir (AG7088) exerts broad-spectrum antiviral activity against enteroviruses and rhinoviruses ^15^. Some efficacy was shown in human trials upon rhinovirus challenge ^16^, but further development was halted due to lack of efficacy in naturally infected patients^17^. The advantage of 3C inhibitors is the high barrier to resistance ^9,18^. Another related 3C protease inhibitor AG-7404 is like pocapavir being developed for the polio endgame ^19,20^.

Nucleoside analogs are successfully being used to treat infections with HIV, HBV, HCV, and herpesviruses ^21^. 7-deaza-2’*C*-methyladenosine (7DMA) was initially developed as an inhibitor of hepatitis C virus ^22^, but was also shown to have antiviral activity against other +ssRNA viruses, including Zika virus ^23^ and human parechovirus ^24^.

The multifunctional viral protein 2C (among others endowed with ATPase and RNA helicase activity) is another attractive target for enterovirus drug development. It is also highly conserved among many enterovirus species. Several 2C targeting molecules have been reported ^25–28^ and one of them – fluoxetine - has been used in a clinical case of chronic enteroviral encephalitis, which led to stabilization and improvement of patient’s condition ^29^. The activity of the 2C-targeting inhibitor SMSK_0213 against EV-71 was discovered in a screening of small molecules in our laboratory (publication in preparation).

Development of resistance to capsid binders is a major problem in the clinical development of this class of compounds. We here explore whether combinations of capsid binder pirodavir with another inhibitor with a different mechanism of action can delay or prevent the development of resistance.

## Results and discussion

### Combined antiviral activity in short term assay

The combined anti-EV-71 activity of pirodavir with either rupintrivir, SMSK_0213 or 7DMA was assessed in a checkerboard format with 8 dilutions of either compound. The inhibition of virus-induced CPE by each compound alone (Table 1) or in combination were determined and the data were analyzed with the SynergyFinder webtool (Table 2) ^30^. The combinations (Table 2 and Fig.1) used in this study resulted in synergy scores of 0.275 (pirodavir and rupintrivir); −1.37 (pirodavir and SMSK) and −2.377 (pirodavir and 7DMA). Overall, over a large surface the combined antiviral effects were rather additive. In certain concentration ranges, the combinations had a trend towards either antagonism or synergy. No cytotoxicity was observed for the tested combinations. The combined effect of capsid binders and rupintrivir was previously shown to be additive ^31^. A synergistic effect of the combination of another 3C inhibitor AG-7404 with two capsid binders was also observed ^19^. This particular combination is further developed as part of the poliovirus eradication program (Antivirals – GPEI).

**Table 1.**
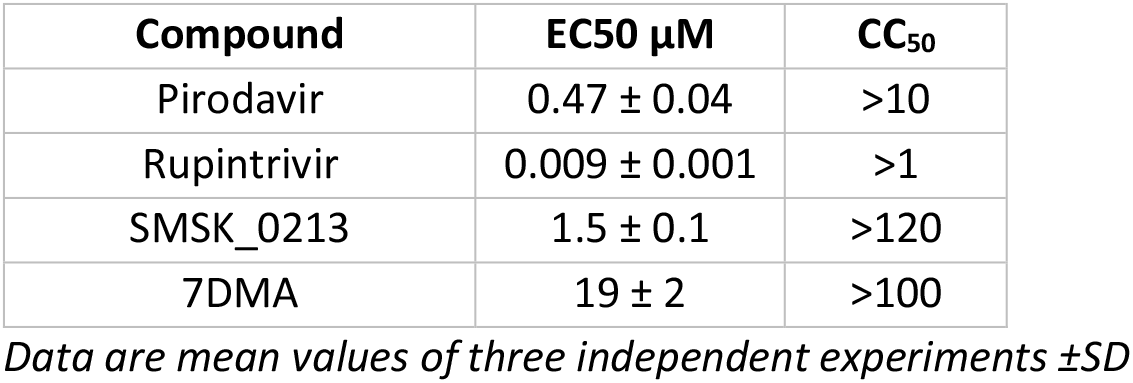
Antiviral activity and cytotoxicity of inhibitors against EV-71.

**Table 2.**
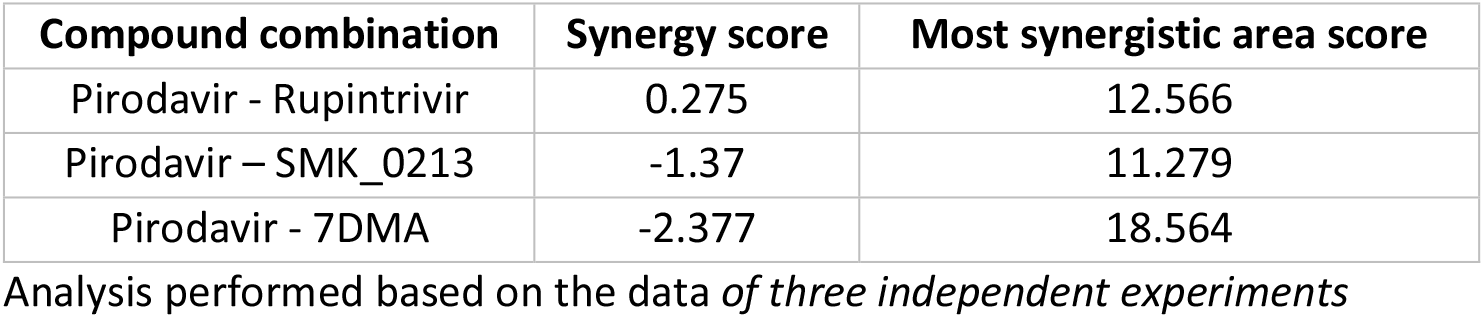
Analysis of the interaction of compounds with the ZIP method.

**Figure 1.**
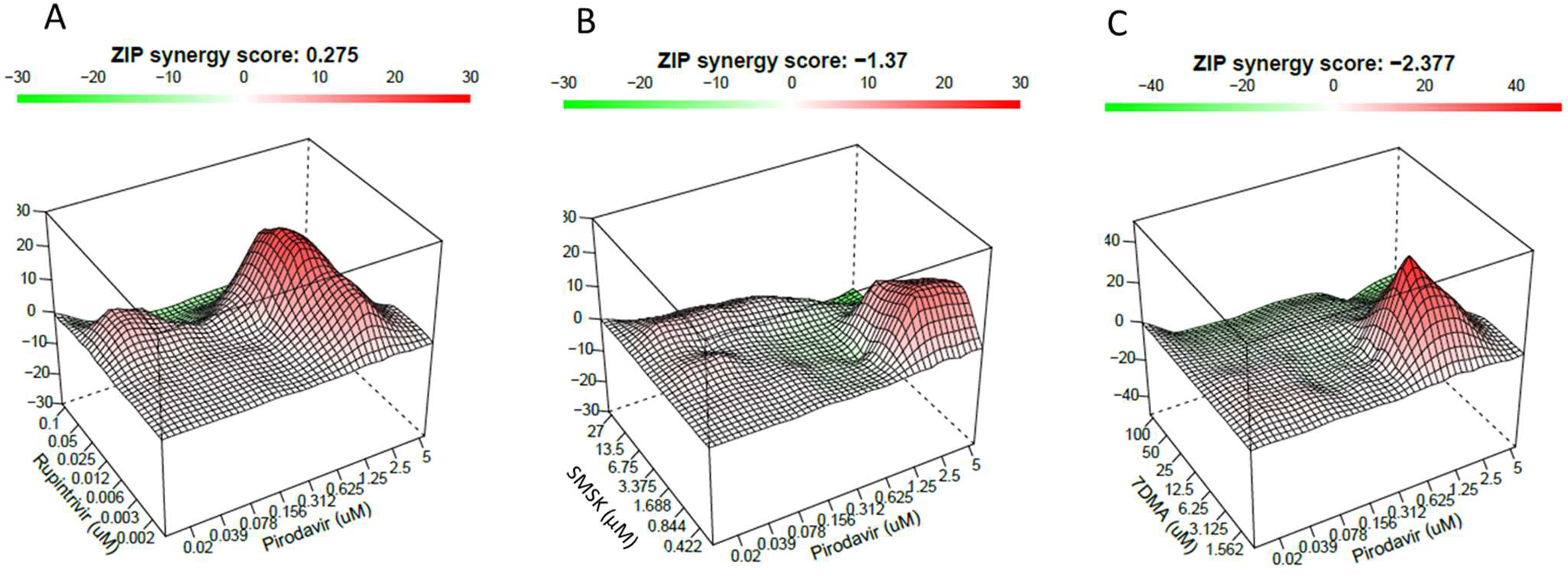
Combined effect of inhibitors on EV-71 infection. Serial dilutions of compounds were added to 96-well plates and the cultures were infected with EV-71. The interaction surface for combinations of (A) pirodavir and rupintrivir, (B) pirodavir and SMSK, (C) pirodavir and 7DMA. The graphics represent the mean of 3 independent experiments.

### Resistance selection in combinations

To assess the effect of combinations on resistance development, serial passaging of EV-71 in combined fixed drug concentrations was performed. Conditions of low antiviral pressure were chosen to still allow for some replication of the virus in such combination of inhibitors was used. All compounds were used at their EC50 concentration (Table 1). For SMSK_0213, a 2xEC50 concentration was used as well. Cultures of RD cells with culture medium supplemented with the respective molecules or combinations thereof were initially infected with EV-71 at a MOI of 0.1. Virus infection was maintained for 3 days, after which the viruses were passaged in the same selection conditions at a 1/1000 dilution. Single treatment and no treatment controls were passaged in parallel. Eight passages were performed and the possible development of resistance under each condition was evaluated in CPE-reduction antiviral assays (Figure 2).

**Figure 2.**
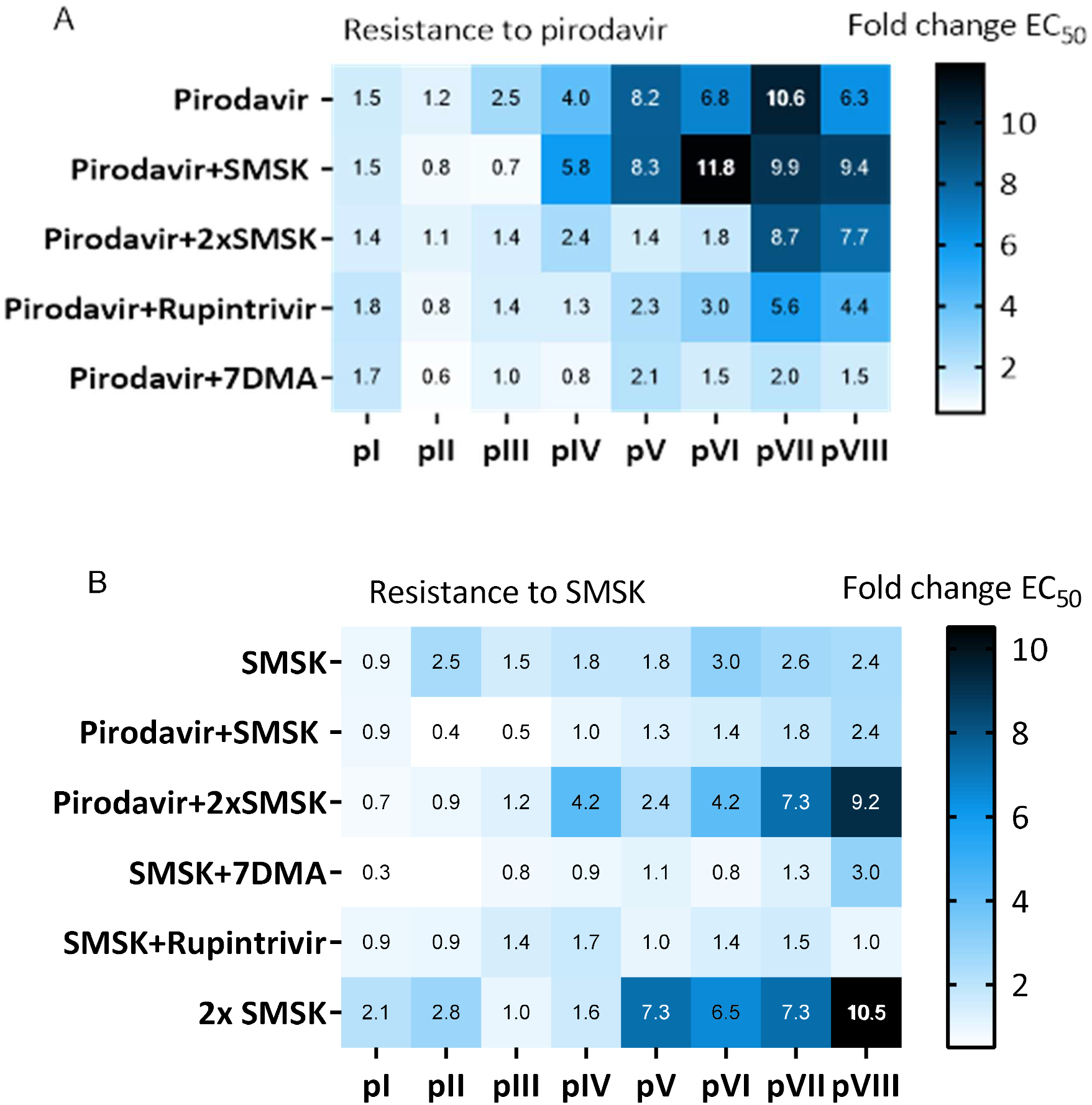
Heatmap of phenotypic resistance development to (A) pirodavir and (B) SMSK. Darker colour indicates a stronger resistant phenotype. The numbers represent fold change in EC_50_ compared to the WT virus based on two independent selection experiments.

### Resistance development to pirodavir

When passaged in the presence of 0.5 μM pirodavir, the virus population started to show a resistant phenotype at passage 3 (2.5-fold change in susceptibility to pirodavir) with further reduction in susceptibility in the subsequent passages. When the 2C-inhibitor SMSK was added as the second selective agent at a fixed concentration of 1.4 μM, a similar pattern of resistance to pirodavir was observed. However, when the concentration of SMSK was doubled to 2.8 μM, a strong resistant phenotype to pirodavir (8.7-fold) was first observed at passage 7. When pirodavir was combined with 0.009 μM rupintrivir emergence of pirodavir resistance was delayed to passage 5. Finally, when pirodavir was combined with the nucleoside analog 7DMA, appearance of pirodavir-resistant variants was entirely prevented until the end of the experiment (8 passages).

### Resistance development to non-capsid binders

Under the conditions used in this study, no phenotypic resistance to either rupintrivir or 7DMA (in monotherapy or in combination with pirodavir) was noted. This is in line with previous reports, where it has been shown that rupintrivir-resistant variant arise at much lower frequency than capsid binder-resistant variants ^9,32^. However, resistance to SMSK was observed (Figure 2). When using 1.4 μM of SMSK, only up to 3-fold resistant populations were selected alone or in combination with pirodavir, 7DMA or rupintrivir. Obviously, 1xEC50 of SMSK is a too low antiviral pressure to select for a resistant population in this experimental setup. At 2xEC50 of SMSK, a high resistance profile after passage 5 was observed in the monoselection with SMSK. In combination with pirodavir, the susceptibility to SMSK decreased (starting from passage 4) and reached a similar high resistant phenotype at passage 7.

### Identification of mutations in double-resistant populations

We next wanted to identify changes in the genomes of the double-resistant virus populations. To that end, deep sequencing of pirodavir+SMSK-resistant populations at the stage of no resistance and full resistance (passages 2 or 5 and 7, respectively) and the input virus was performed (Table 3). Mutations S196P in VP1 (observed in the pirodavir-only selected population) and I113V (identified in a double-resistant population) are associated with the resistance to pirodavir: the reverse-engineered virus with the S196P substitution had a reduced susceptibility to pirodavir (EC50 of 3.2±1.5 μM, 7-fold change over WT) and the introduction of I113V mutation increased the EC50 of pirodavir to 10±0.4 μM (21-fold change over WT). The I113 residue has been shown to interact with another capsid binder, WIN51711, whereas the S196 residue is located in the drug-binding pocket next to M195 which is also involved in drug interaction ^33^. Mutation T173M in 2C conferred resistance to SMSK with an EC50 of 5.7±0.3 μM (~4-fold change over WT) when reversed-engineered into the WT virus backbone. Viruses resistant to 2C inhibitors have been reported for several compounds (TBZE-029, HBB, Guanidine HCl, hydantoin, fluoxetine) ^26–28,34–36^. Some of these resistant viruses have an attenuated virus growth ^28^ or even a compound-dependent phenotype ^27,37^. We observed slower development of CPE in cultures infected with the 2C T173M variant, which may explain the low frequency of this variant in the double-resistant population (17.7-24.7%). The low frequencies of both the VP1 I113V and T173M variants might also be due to the low compound pressure used in this study, which might not be enough to let the resistant population overgrow the wild type. However, the resistant phenotype was still detectable in a CPE-reduction antiviral assay, where the compound pressure is higher than during passaging.

**Table 3.**
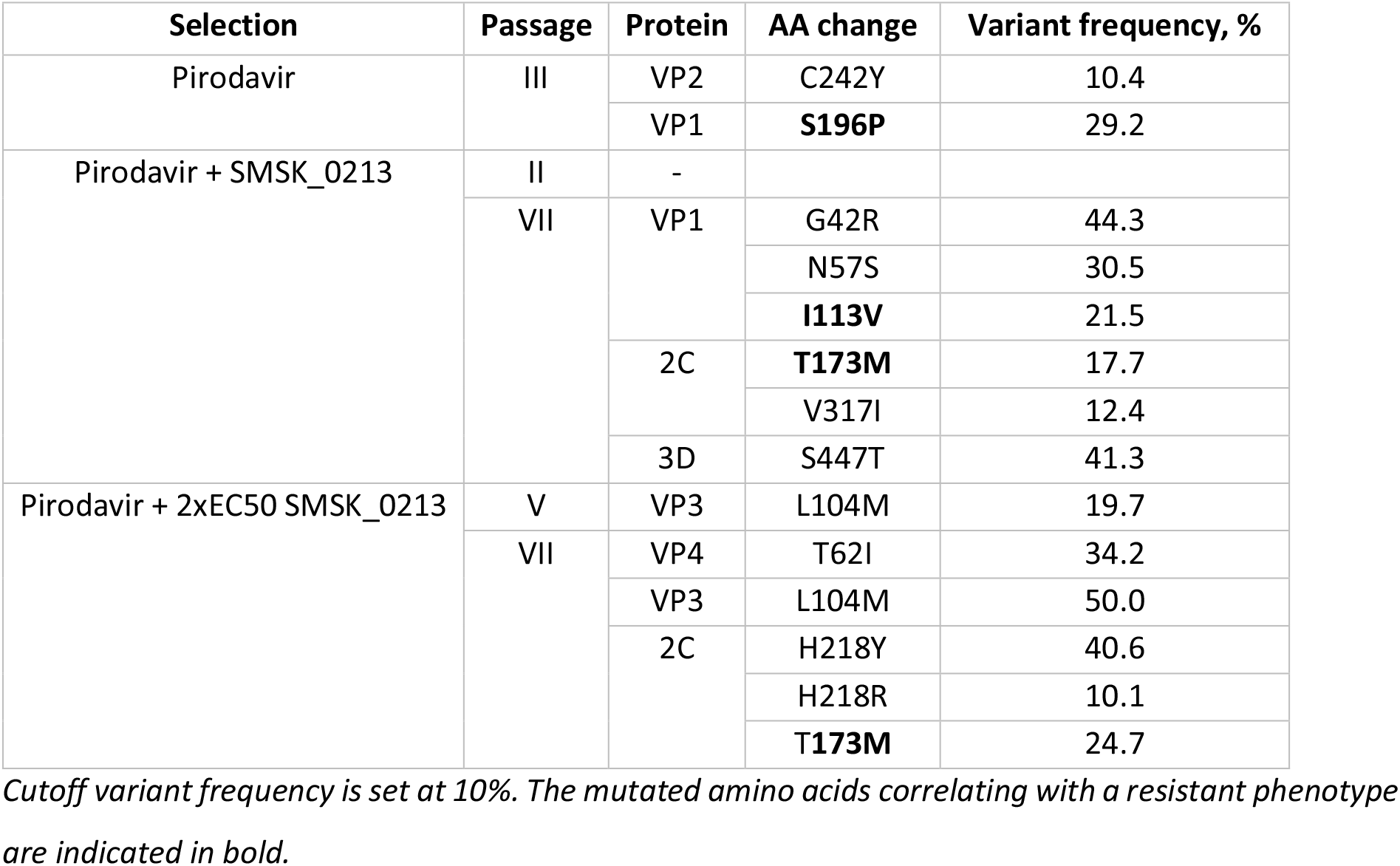
Amino acid substitutions identified with deep sequencing of double-resistant populations.

## Conclusion

The use of combinations of antiviral compounds is common in the treatment of infections with HIV and HCV. The GPEI considers the development of a combination of two antiviral molecules as an essential tool in the polio endgame. Monotherapy with the capsid binder pocapavir resulted rapidly in the selection of drug-resistant variants in healthy volunteers that had been experimentally infected with OPV ^12^. Several studies have explored the combined effect of antiviral compounds on enterovirus infections ^19,31,38,39^. We reasoned however that although it might be of interest to identify molecules that may result in a synergistic antiviral effect (or exclude those that would result in an antagonistic effect), it would be even more relevant to understand which combinations may have the potential to delay or even prevent resistance development to either one or both molecules. To this end, we studied the potential development of (single or double) resistance to either combinations of a capsid binder (pirodavir) with either a protease inhibitor, a nucleoside polymerase inhibitor or a 2C targeting molecule. We here report a delay in resistance development to pirodavir in presence of even a low concentration of rupintrivir. The nucleotide polymerase inhibitor 7DMA even prevented the appearance of pirodavir-resistant variants during the time-span of the experiment. On the other hand, when combined with the 2C-targeting molecule SMSK, double-resistant variants emerged, although slightly later than in monotherapy. Of note, no strong antagonistic effects were observed in any of the combinations tested, indicating that combinations of capsid binders (or at least pirodavir) with a 3C protease inhibitor or a nucleotide inhibitor may be considered. Obviously, studies as presented here will have to be repeated with future drug-candidates.

Combinations of enterovirus inhibitors with different mechanism of actions (in this study entry and replication inhibitors) can delay or even prevent the development of drug-resistant variants at least in vitro. Studies reported here may be instrumental to circumvent the challenge of rapid emergence of resistance to capsid binders in vivo. There is a preponderance of knowledge on drug-combination for the treatment of HIV and HCV. Several combinations are highly efficient and avoid the development of resistant variants. Combinations of enterovirus inhibitors may have to be designed as a tool in the polio endgame but may as well be needed for the treatment of some of the life-threatening enterovirus infections.

## Acknowledgements

This project has received funding from the European Union’s Horizon 2020 research and innovation programme under the Marie Sklodowska-Curie grant agreement No 642434.

