## Supplementary material for "Assessing the *in vitro* resistance development in Enterovirus 71 in the context of combination antiviral treatment": Materials and methods

### Cells and viruses

Human Caucasian embryo rhabdomyosarcoma (RD, ECACC 85111502) were cultured at 37°C in 5% $CO_2$ incubators in MEM Rega3 medium (Gibco) supplemented with 10% FBS, 2mM L-glutamine (Gibco) and 0.075% sodium bicarbonate (Gibco). For infection experiments 2% medium was used.

Enterovirus A71 BrCr strain was a kind gift from Prof. F. van Kuppeveld (University of Utrecht, The Netherlands). The virus stock was grown in RD cells in 2% FBS MEM Rega3 medium.

### Compounds

Pirodavir was synthesized by Prof. G. Pürstinger (University of Innsbruck). Rupintrivir was provided by Axon Medchem (The Netherlands). SMSK_0213 was synthesized by Prof. S. Mikhailov (Engelhardt Institute of Molecular Biology, Moscow, Russia). 7-deaza-2'-*C*-methyl-D-adenosine (7DMA) was purchased from Carbosynth (Berkshire, UK). Compound stocks were prepared in DMSO at a concentration of 10 mM and diluted in cell culture medium for experimental use.

### Antiviral CPE-reduction assays

Cells were seeded in 96-well plates ($20*10^3$ cells/well) in 2% FBS MEM Rega3 medium and kept overnight. The next day serial dilutions of the compounds were added to the cells. These cultures were then infected with EV-71 at a MOI 0,01 and kept for 3 days. Microscopic evaluation of cytopathic effects and cell viability readout with MTS reagent [3-(4,5-dimethylthiazol-2-yl)-5-(3-carboxymethoxyphenol)-2-(4-sulfophenyl)-2H-tetrazolium, inner salt; Promega] were performed for assessing the susceptibility of EV-71 isolates to the antiviral effect of compounds.

### Deep sequencing and data analysis

An optimized sample preparation protocol for viral metagenomics—NetoVIR[1], was used for the analyses of the different mutant viruses. Briefly, cell cultures were centrifuged (17,000 g for 3 min), and 150 µl supernatant was used for filtration (0.8 µm pore size) to enrich for viral particles. The filtrate was then treated with a cocktail of Benzonase (Novagen) and Micrococcal Nuclease (New England Biolabs) in a homemade buffer (1 M Tris, 100 mM $CaCl_2$, and 30 mM $MgCl_2$) to digest free-floating nucleic acids. DNA and RNA were extracted (QIAGEN Viral RNA mini kit), reverse-transcribed, and randomly amplified using a slightly modified Whole Transcriptome Amplification 2 (WTA2) Kit procedure (Sigma-Aldrich). WTA2 products were purified, and the libraries were prepared for Illumina sequencing using the NexteraXT Library Preparation Kit (Illumina). A cleanup after library synthesis was performed using a 1.8 ratio of Agencourt AMPure XP beads (Beckman Coulter, Inc.). Sequencing of the samples was performed on a NextSeq500 High throughput platform (Illumina) for 300 cycles (2 × 150 bp paired ends).

All the reads were filtered based on their quality before aligning them to the reference genome. The quality control analysis was performed using FASTQC [1]. Based on the quality control analysis the reads were processed to trim low quality bases and remove the adapter sequences using Trimmomatic [2].

Next the pre-processed reads were aligned to the reference genome (in our case a consensus sequence ie the largest scaffold/contig) to map back the reads to their respective positions using Bowtie2 [3]. The alignment output file (.sam file) contains the position of all the mapped reads which were used for further analysis. This involves conversion of the .sam file to a compressed .bam format file, sorting and indexing of the .bam file using samtools [4].

Finally the processed alignment output was used to scan for possible variants using freebayes, a haplotype-based variant detector [5]. A variant allele frequency of 0.1 was used as threshold. The output of the variant calling process is a VCF formal file, which comprises of the following fields; position of the variant, reference base(s) present on genome, alternate base(s) present on genome, variant allele frequency (VAF), reference allele frequency (RAF) and the depth.

### Site-directed mutagenesis

Mutagenesis was performed on the plasmid pT7/EV-A71BrCr[6] by using the QuikChange II XL Site-Directed Mutagenesis Kit (Agilent Technologies). RNA was obtained by using a T7 RiboMAX Large Scale RNA Production System (Promega) and infectious viruses were generated by transfecting RNA into RD cells with the TransIT-mRNA Transfection Kit (Mirus). Mutations in the genome of engineered viruses were verified by Sanger sequencing.
